# Precision digital mapping of endogenous and induced genomic DNA breaks by INDUCE-seq

**DOI:** 10.1101/2020.08.25.266239

**Authors:** Felix M Dobbs, Patrick van Eijk, Mick D Fellows, Luisa Loiacono, Roberto Nitsch, Simon H. Reed

## Abstract

Understanding how breaks form and are repaired in the genome depends on the accurate measurement of the frequency and position of DNA double strand breaks (DSBs). This is crucial for identification of a chemical’s DNA damage potential and for safe development of therapies, including genome editing technologies. Current DSB sequencing methods suffer from high background levels, the inability to accurately measure low frequency endogenous breaks and high sequencing costs. Here we describe INDUCE-seq, which overcomes these problems, detecting simultaneously the presence of low-level endogenous DSBs caused by physiological processes, and higher-level recurrent breaks induced by restriction enzymes or CRISPR-Cas nucleases. INDUCE-seq exploits an innovative NGS flow cell enrichment method, permitting the digital detection of breaks. It can therefore be used to determine the mechanism of DSB repair and to facilitate safe development of therapeutic genome editing. We discuss how the method can be adapted to detect other genomic features.

---

DNA double-strand breaks (DSBs) are the most toxic of all DNA lesions, directly compromising genome stability. Other than causing cell death, failure to repair DSBs accurately can lead to a range of structural genomic alterations associated with carcinogenesis^1–3^. Low-level physiological breaks can arise sporadically because of normal cellular processes such as DNA replication, transcription, and chromatin looping^2^. DSBs are also induced at high frequencies as programmed events in specialised cell-types: DSBs occur as natural intermediates of the DNA repair mechanisms that allow for antigen receptor gene recombination in developing and antigen-stimulated lymphocytes; intentionally induced DSBs also represent the initiating substrates of meiotic recombination reactions that enable genetic diversification in the germline^4^. A variety of exogenous physical and chemical agents, such as ionising radiation, chemotherapeutic drugs and more recently, CRISPR genome editing technologies^5, 6^, are also potent inducers of DSBs.

Until now, precise measurement of the full complement of genomic DSBs present in cells has not been possible. This is because most physiological breaks are infrequent and occur stochastically throughout the genome. Existing methodologies typically measure DSBs reliably only when they exist at recurrent ‘hotspots’, or when they are induced at defined genomic locations by sequence-directed nucleases. These next generation sequencing (NGS) based methods fall broadly into three categories: indirect break labelling using proteins as a proxy for breaks (e.g γH2AX ChIP-seq, DISCOVER-seq)^7, 8^; *indirect* labelling of repaired breaks (e.g. GUIDE-seq, HTGTS)^9, 10^ and finally, *direct* labelling of unrepaired break-ends in cells (e.g. BLESS, DSBCapture, END-seq, BLISS)^11–13^. All current methodologies suffer from distorted break measurements caused by PCR amplification bias during the standard DNA library preparation required for NGS-based break sequencing^14^. PCR amplification distorts the representation of the actual distribution of DSBs present in the genome. This well-known phenomenon makes the quantification of genomic DSBs impossible^15–17^. For most NGS applications, this nuance is not a significant problem. But for the quantitative, genome-wide measurement of specific features, such as DSBs, PCR-amplification bias introduces high levels of noise into a system where the signals (genomic DSBs) are already very low. To overcome the attenuation of the DSB signal by the noise associated with PCR amplification, we designed a novel DNA library preparation protocol that avoids amplification and improves the DSB signal by break-sequence enrichment *directly* on the Illumina flow cell. By improving the signal-to-noise ratio for DSB detection by noise filtering, instead of signal amplification, we obtain a digital measurement of genomic breaks, where one sequence read is equivalent to one labelled DSB-end in the genome. Here, we describe the novel method INDUCE-seq for the direct, digital measurement of genomic DSBs. We demonstrate its ability to locate and characterise endogenous and induced DSBs caused by different nucleases including CRISPR/Cas9.

Break measurement by INDUCE-seq is achieved via a two-stage, PCR-free library preparation (**Fig. 1 and Supplementary Fig. 1a**). Stage one consists of labelling *in situ* end-prepared DSBs via ligation of a full-length, chemically modified P5 adapter. In stage two, extracted, fragmented, and end-prepared gDNA is ligated using a second chemically modified, half-functional P7 adapter. The resulting DSB-labelled DNA fragments that comprise both the P5 and half-functional P7 adapters can hybridise with the Illumina flow cell and subsequently be sequenced using single-read sequencing. DNA fragments that do not possess the P5 adapter remain nonfunctional, since they lack the sequence required to hybridise to the flow cell. This methodology enables the enrichment of functionally labelled DSB sequences and the elimination of all other genomic DNA fragments, which would otherwise contribute to system noise. The avoidance of break-sequence amplification produces a sequencing output where a *single* sequencing read is equivalent to a *single* labelled DSB end in the cell (compare **Supplementary Fig. 1a and 1b**, see **Supplementary Fig. 2a**). This important innovation generates a digital DNA break signal, enabling the direct detection, and therefore quantification, of genomic DSBs by sequencing.

**Fig 1:**
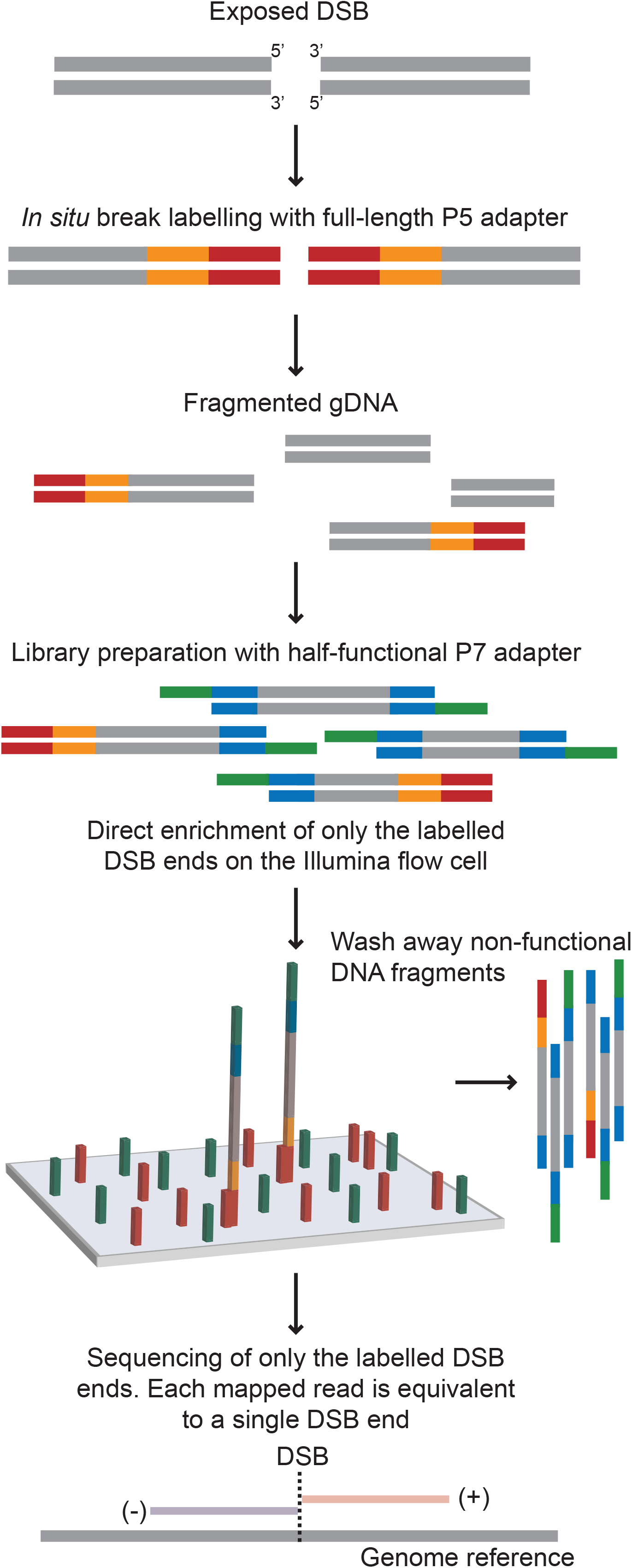
The INDUCE-seq workflow. *In situ* break labelling in fixed and permeabilised cells is performed by ligating a full-length chemically modified P5 sequencing adapter to end-prepared DSBs. Genomic DNA is then extracted, fragmented, end-prepared and ligated using a chemically modified half-functional P7 adapter. Resulting DNA libraries contain a mixture of functional DSB-labelled fragments (P5:P7) and nonfunctional genomic DNA fragments (P7:P7). Subsequent Illlumina sequencing of INDUCE-seq libraries enriches for DNA-labelled fragments and eliminates all other non-functional DNA. As the INDUCE-seq library preparation is PCR-free, each sequencing read obtained is equivalent to a single labelled DSB end from a cell.

Following *in situ* break labelling, currently available DSB detection methods BLISS, DSBCapture and END-seq, all employ an enrichment protocol to separate DSB-labelled DNA fragments from the remaining genomic DNA. This is followed by a PCR-based library preparation, and Illumina sequencing (**Supplementary Fig. 1b**). This results in a readout where a single read is not equivalent to a single break. Therefore PCR error-correction, such as a unique molecular identifier (UMI), is required to attempt DSB quantification (**Supplementary Fig. 1b** and **Supplementary Fig. 2b**)^18^. Importantly, the novel INDUCE-seq is compatible with any of the *in situ* DSB labelling protocols reported to date.

To demonstrate the performance of the INDUCE-seq, we first examined the genome-wide DSB distribution induced by a high-fidelity HindIII restriction endonuclease. This approach has been used to benchmark BLISS, END-seq and DSBCapture^12, 13, 18^. As shown in **Fig. 2a**, INDUCE-seq simultaneously detects hundreds of millions of highly recurrent HindIII-induced DSBs, in addition to hundreds of thousands of lower-level endogenous DSBs from within the same sample. This enables the precise quantification and characterisation of endogenous DSBs in the genome for the first time. **Supplementary Fig. 3** shows two examples of endogenous DSB detection using INDUCE-seq in HEK293 cells. Collectively, these observations show the remarkably broad dynamic range achievable with INDUCE-seq. **Fig 2.b** shows that INDUCE-seq detects the expected HindIII cleavage pattern, where two semi-overlapping symmetrical blocks of sequencing reads map to the known HindIII cleavage positions on both strands. Thus, INDUCE-seq can be used to precisely measure DSB-end structures at single-nucleotide resolution. We measured a dramatic increase in breaks-per-cell following treatment of cells with HindIII; from fewer than 10 endogenous breaks per untreated cell, to more than 3,000 enzyme-induced breaks per treated cell. This demonstrates that INDUCE-seq is capable of quantitatively measuring breaks-per-cell across three orders of magnitude (**Fig. 2c**). Compared with an equivalent experiment using DSBCapture to detect EcoRV induced breaks^12^, we found that a greater proportion of INDUCE-seq reads were mapped to restriction sites (**Fig. 2d**). 96.7% of aligned reads were mapped to restriction sites, representing a 25% improvement in fidelity of break detection over DSBCapture. Significantly, INDUCE-seq uses 800-fold fewer cells than DSBCapture, whilst identifying a similar proportion of HindIII restriction sites (92.7%) to that identified by DSBCapture for EcoRV (93.7%) (**Fig. 2e**). In addition to on-target HindIII sites, we also identified a substantial number of DSBs at a variety of HindIII off-target sequences; cryptic sites that differ by one or two mismatching bases (**Fig. 2f**). The total number of HindIII-induced DSBs measured by INDUCE-seq ranged from ~150,000,000 across ~780,000 on-target sites to just five DSBs at the lowest ranking off-target site (**Fig. 2f**). INDUCE-seq therefore quantitatively detects breaks across seven orders of magnitude, vastly enhancing the dynamic range of break detection over current methods. Detection of breaks at AsiSI sites in live DiVA cells was also enhanced. INDUCE-seq detected the presence of breaks at 230 AsiSI sites despite sequencing 40-fold fewer reads than a corresponding DSBcapture experiment, and 23-fold fewer reads than BLISS^12, 19^. This represents an increase over the 214 sites detected by BLISS, and 121 by DSBCapture. Therefore INDUCE-seq is significantly more sensitive, efficient, and cost effective than current methods (**Fig. 2g**).

**Fig 2:**
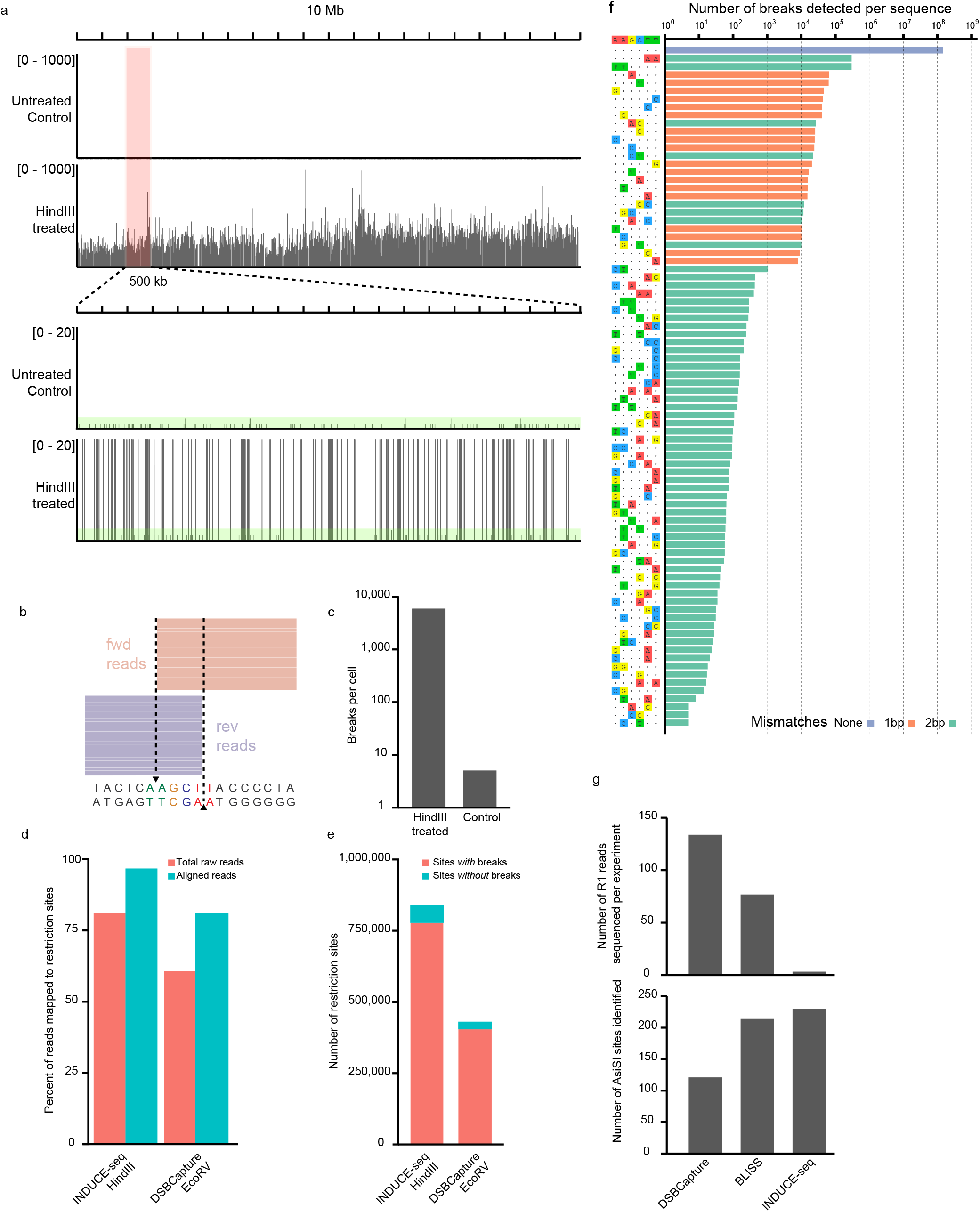
INDUCE-seq demonstrates unparalleled sensitivity and dynamic range when compared to alternative DSB sequencing technologies. (**a**) INDUCE-seq detects both highly recurrent induced DSBs and low-level endogenous DSBs simultaneously with high resolution. Genome browser view (IGV) of INDUCE-seq reads mapped to a 10Mb section of the genome from HEK293T cells following *in situ* cleavage with the restriction endonuclease HindIII. (**Top panel**) Highly recurrent enzyme-induced breaks represent the vast majority of reads when viewed at low resolution (10Mb, 0-1000 reads). (**Bottom panel**) A high resolution view (pink highlight, 500kb, 0-20 reads) reveals low level single endogenous breaks present in both the untreated sample and amongst the recurrent HindIII-induced breaks (Green highlight). (**b**) Mapping of INDUCE-seq reads at a HindIII target site demonstrates precision of single-nucleotide break mapping on both sides of the break. (**c**) Quantification of breaks measured per cell for the HindIII treated and control samples. INDUCE-seq quantitatively detects breaks-per-cell across 3 orders of magnitude between samples. (**d and e**) Comparison between INDUCE-seq and DSBCapture in detecting *in vitro* cleaved restriction sites by the enzymes HindIII and EcoRV. (**d**) A greater proportion of reads sequenced and aligned to the genome were mapped to restriction sites using INDUCE-seq. (**e**) Using 800-fold fewer cells, INDUCE-seq identifies a similar proportion of HindIII restriction sites (92.7%) to that identified by DSBCapture for EcoRV (93.7%). (**f**) The dynamic range of induced DSB detection using INDUCE-seq. In addition to breaks identified at HindIII on-target sequences (AAGCTT), multiple 1bp and 2bp mismatching off-target sites were also identified. INDUCE-seq measured induced break events spanning 8 orders of magnitude, from ~150 million breaks identified at HindIII on-target sites, to 5 breaks identified at the least frequent off-target. (**g**) Comparison between INDUCE-seq, DSBCapture and BLISS in detecting AsiSI induced breaks in live DiVA cells. The number of reads sequenced (**top panel**) is compared to the number of AsiSI sites identified for each experiment (**bottom panel**). INDUCE-seq detects the greatest number of AsiSI sites using 40-fold fewer reads than DSBCaptue and 23-fold fewer reads than BLISS.

Having established the characteristics of break detection by INDUCE-seq, next we applied it to the detection of CRISPR/Cas9-induced on and off-target DSBs in the genome. This analysis is of central importance in safety profiling for the development of CRISPR-based therapies. Following RNP nucleofection of HEK293 cells with the extensively characterised EMX1 sgRNA, DSBs were measured at 0, 7, 12, 24 and 30 hours post-nucleofection, and off-targets were defined by a custom data analysis pipeline (**Fig. 3a and Supplementary Data 1**). This off-target discovery pipeline is described in **Supplementary Fig. 4**. **Supplementary Fig. 5** shows the effect on true and false positive off-target discovery, following the full spectrum of filtering parameters used in the workflow. Finally, **Supplementary Fig 6a** demonstrates the statistical significance of off-target detection in relation to different filter conditions. This analysis could be used to select the appropriate filtering threshold for off-target discovery with any given sgRNA. For EMX1, filter condition 7 (**Supplementary Fig 6a**, highlighted red) was selected for subsequent analysis as shown in **Fig. 3a**, which reports the break number discovered at the on-target and each of the 60 off-targets detected.

**Fig 3:**
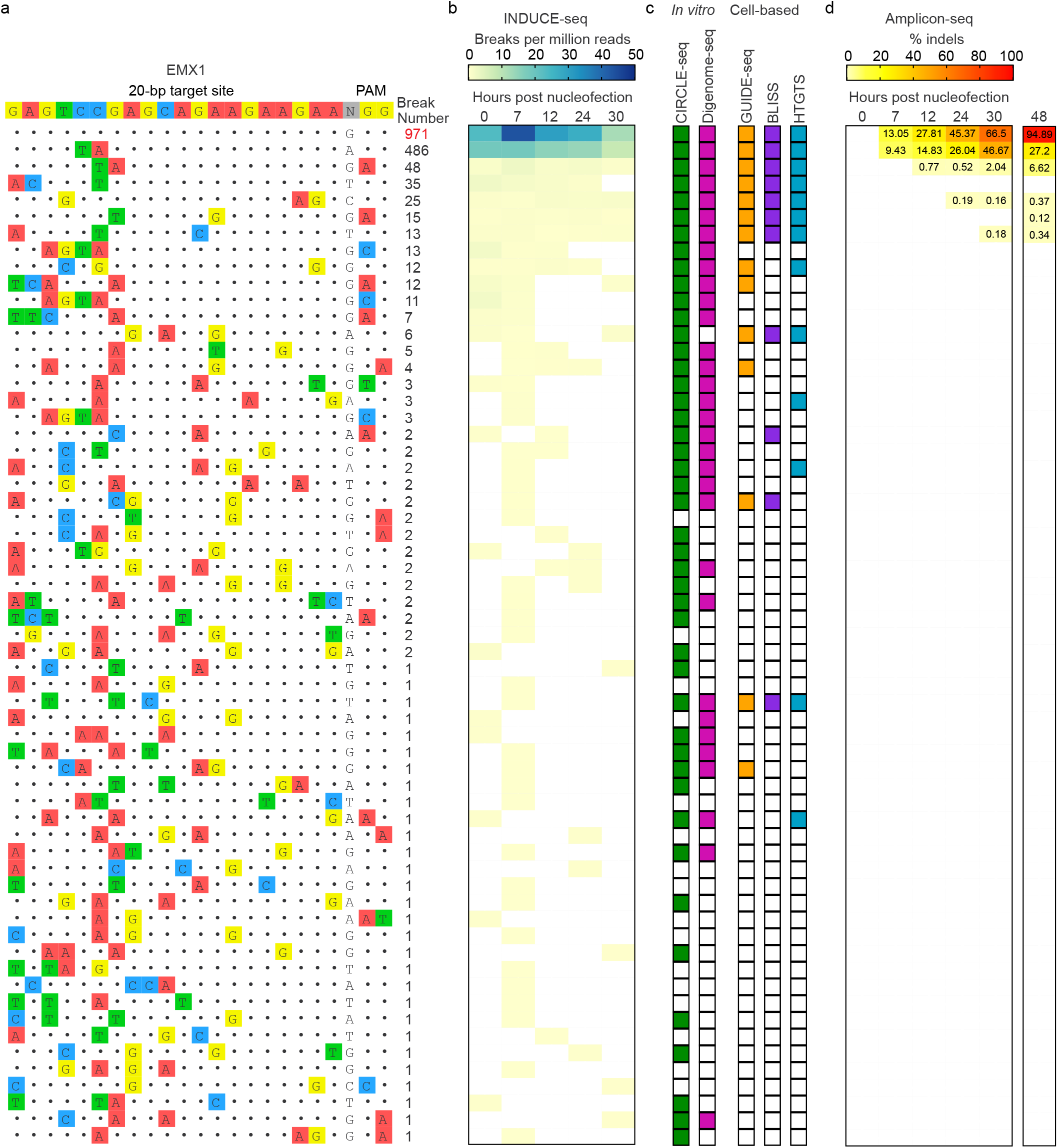
INDUCE-seq sensitively discovers and quantifies CRISPR/Cas9 induced on- and off-target DSBs. (**a**) On- and multiple off-target sequences and the number of breaks identified using INDUCE-seq for the EMX1 sgRNA. Mismatching bases from the target sequence are highlighted and colour coded. (**b**) INDUCE-seq reveals the kinetics of EMX1-induced DSB formation in a cell population during the editing process. Quantification of the number of breaks detected per million reads for each sample revealed high Cas9 activity both on- and off-target immediately following cell nucleofection. (**c**) The comparison between off-targets identified by INDUCE-seq with established *in vitro* methods CIRCLE-seq and Digenome-seq, in addition to cell-based methods GUIDE-seq, BLISS, and HTGTS. INDUCE-seq detects many off-targets that were previously only identifiable by *in vitro* approaches. Substantially more off-target sites were identified than by any of the current cell-based methods. INDUCE-seq also identifies multiple off-targets not detected by any other method. (**d**) Ampliconsequencing to measure the indel frequency at INDUCE-seq identified off-targets. Amplicon sequencing identifies only 4 of the 60 off-targets discovered using INDUCE-seq and is limited by the background indel false-discovery rate of 0.1%. (**d, far-right**) Indel frequency reported previously for EMX1 48 hours post RNP nucleofection.

This experiment reveals a profile of the kinetics of break induction by the EMX1 guide, offering insights into the mechanism of the CRISPR/Cas9 editing process. As shown in **Figure 3b**, the majority of on- and off-target activity was observed immediately following nucleofection and during the early stages of the time course shown. When compared to existing technologies, we find that INDUCE-seq significantly outperforms alternative cell-based methods GUIDE-seq, BLISS and HTGTS, as well as capturing several sites that were previously only identified using *in vitro* off-target discovery methodologies CIRCLE-seq and Digenome-seq (**Fig. 3c**). Importantly, INDUCE-seq also detects novel off-target break sites not previously detected by any other method (**Supplementary Fig. 7**).

Finally, using DNA from the same samples, we measured the editing outcome at INDUCE-seq defined on- and off-target sites, using amplicon sequencing. This method only detects indels above a background false-discovery rate of 0.1%^20–22^. Consequently, evidence of editing was detected at the on-target, and only four of the 60 off-target sites that we discovered (**Fig. 3d**). This observation is in agreement with a previous study using GUIDE-seq for off-target detection^23^, which identified five off-targets with indel frequencies >0.1% at 48 hours post-nucleofection of HEK293 cells with EMX1 RNP (**Fig. 3d, 48h**). Collectively, we show that INDUCE-seq discovers CRISPR-induced off-target DSBs with higher sensitivity than indel detection using amplicon sequencing. This observation highlights the need for more sensitive methods for the detection of genome editing outcomes, to accurately evaluate the safety of genome editing. We note that the sgRNA-specific cleavage pattern observed reflects the editing outcome at the on-target and top two off-target sites (**Supplementary Fig. 8**). This raises the intriguing possibility of using the CRISPR-induced DSB pattern to model and predict the editing outcome.

We developed a novel PCR-free methodology to prepare DNA libraries for next generation sequencing of genomic DSBs. This advance, which exploits enrichment of break sequences using the Illumina flow cell, generates a digital output for measurement of breaks in cells for the first time. Our approach overcomes the problem of poor signal-to-noise ratios for DSB detection associated with PCR-amplification employed in standard NGS library preparations. Our novel INDUCE-seq adapter design permits the sequencing flow cell to be used to enrich for labelled DSB sequences, thereby avoiding the need for their amplification by PCR. Improvement in the signal-to-noise ratio is instead achieved by filtering the ‘noisy’ break ends generated during DNA fragmentation that are not associated with physiological DSBs found in cells. We demonstrate the characteristics of INDUCE-seq for measuring genomic DSBs in a range of different applications. We reveal its capacity to detect sensitively and quantitatively low-level endogenous, as well as high-level restriction enzyme-induced breaks simultaneously. This has not been possible previously without the need for complex error-correction methods that have their own limitations. We compare INDUCE-seq with the currently available break detection methods to demonstrate the improved performance, scalability, ease of use, and cost effectiveness. These are all essential features of an assay that can be used to assess the safety profiling of synthetic guides for CRISPR genome editing. We demonstrate how INDUCE-seq compares to several of the current CRISPR off-target detection methods for the measurement of editing by the EMX1 sgRNA. We reveal that in addition to detecting many off-targets reported previously by five other methods, INDUCE-seq also identifies a significant number of novel off-target sites. INDUCE-seq may be a valuable method for safety profiling and synthetic guide RNA design for the future development of genome editing as a therapeutic modality. Finally, we note that this methodology can be adapted for the detection of a range of other genomic features that can be end-labelled in this way. Such features include genome-wide mutations, single strand breaks and gaps, as well as other types of DNA damage that can be converted into breaks and subsequently ligated using this novel combination of sequencing adapters. The development of INDUCE-seq and its derivative assays could have significant implications in a range of different biomedical applications.

---

**Supplementary Fig 1:**
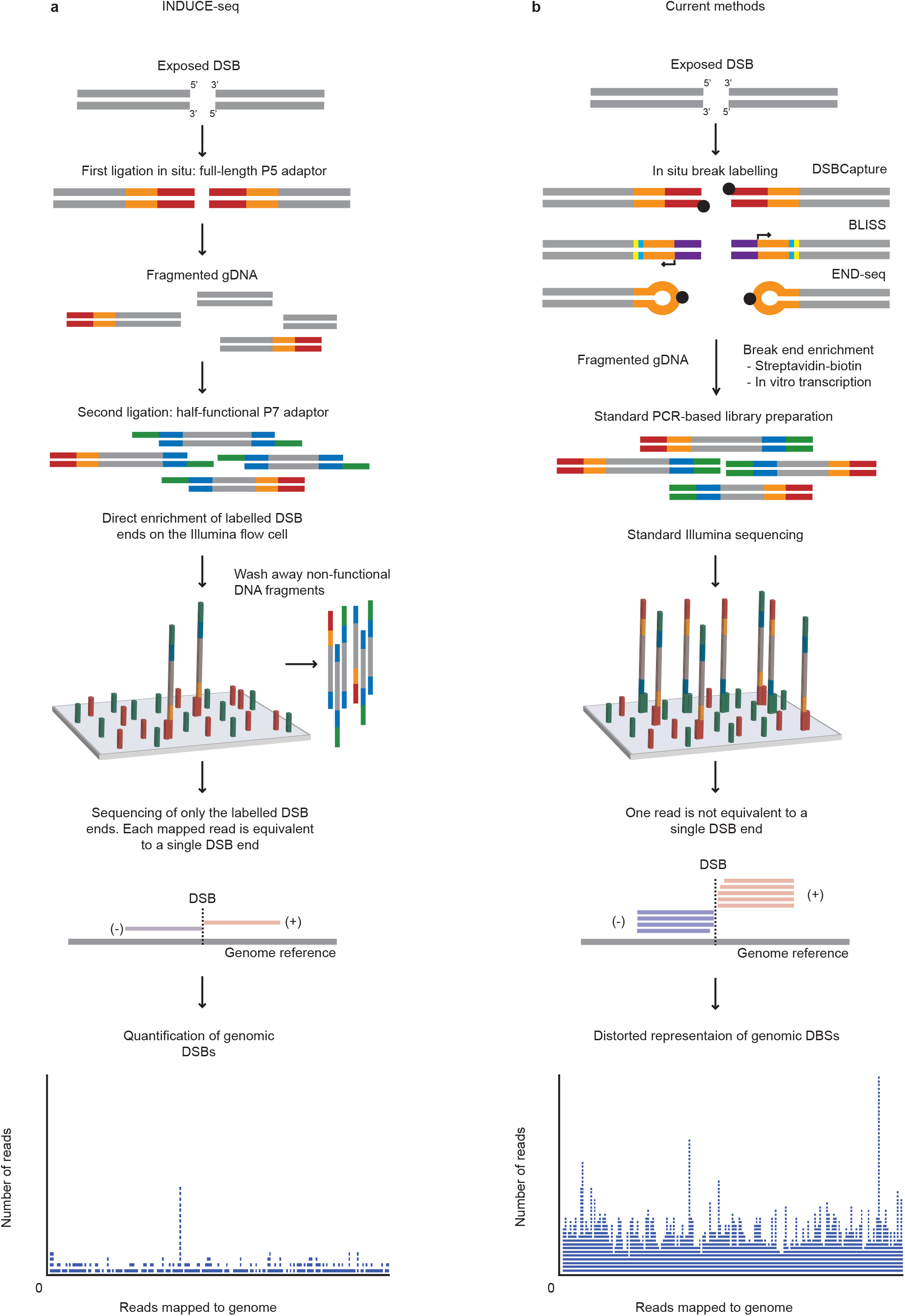
Comparison of INDUCE-seq and current DSB mapping workflows. (**a**) Overview of INDUCE-seq workflow. The sequencing of INDUCE-seq libraries generates a quantitative output where one read is equivalent to one break. (**b**) Overview of the DSBCapture, BLISS and END-seq workflows. Sequencing following standard library construction generates an output where one read is not equivalent to a single DSB.

**Supplementary Fig 2:**
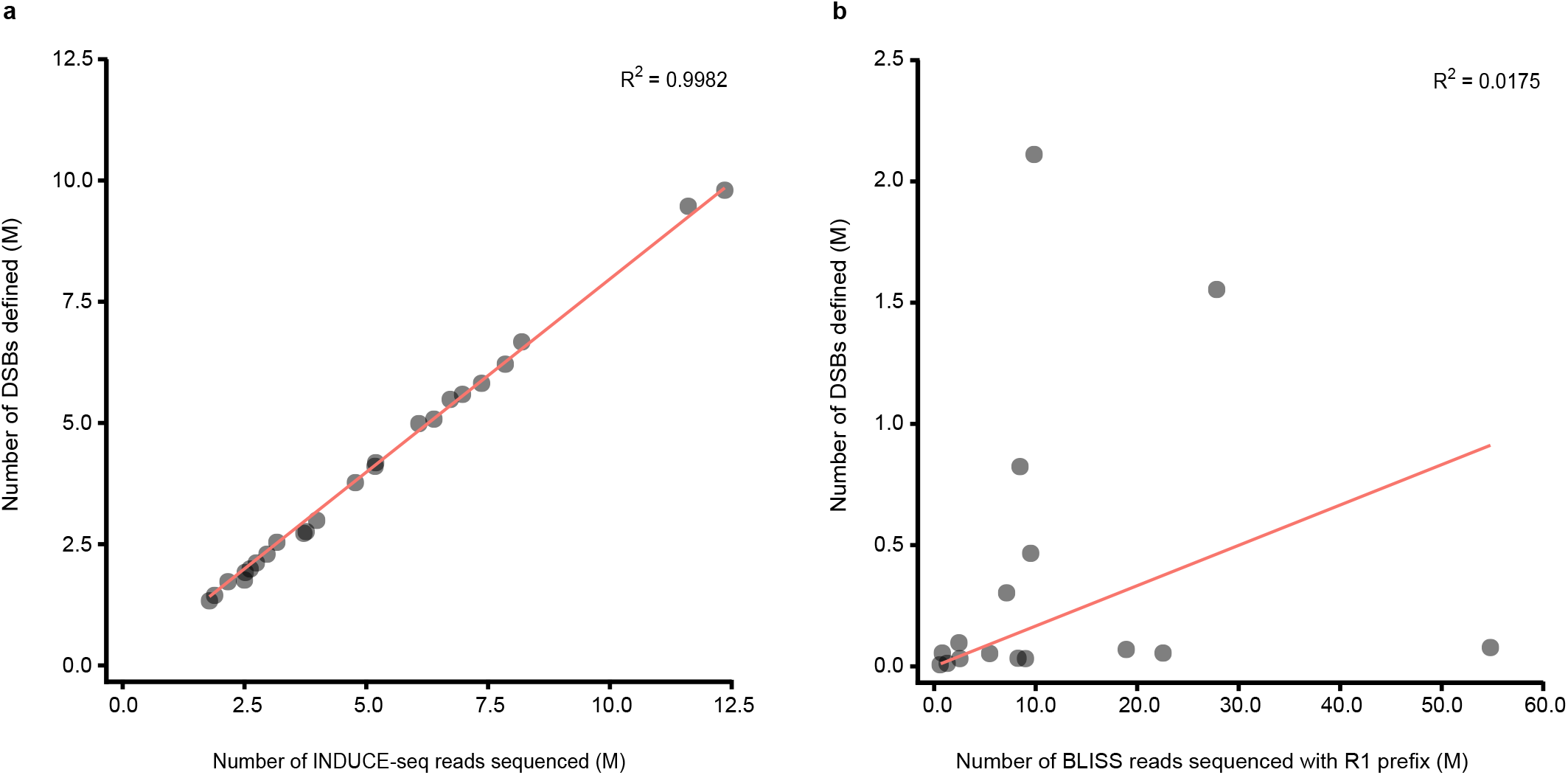
Comparison between the number of reads sequenced and number of DSBs defined for INDUCE-seq and BLISS NGS libraries. (**a**) Scatter plot of the number of INDUCE-seq reads sequenced (millions) and the number of breaks defined (millions) from individual INDUCE-seq experiments. (**b**) Scatter plot showing the number of BLISS reads sequenced (millions) with the correct read 1 (R1) barcode prefix and the number of breaks defined (millions), following duplicate-removal using UMI correction, from individual BLISS experiments.

**Supplementary Fig 3:**
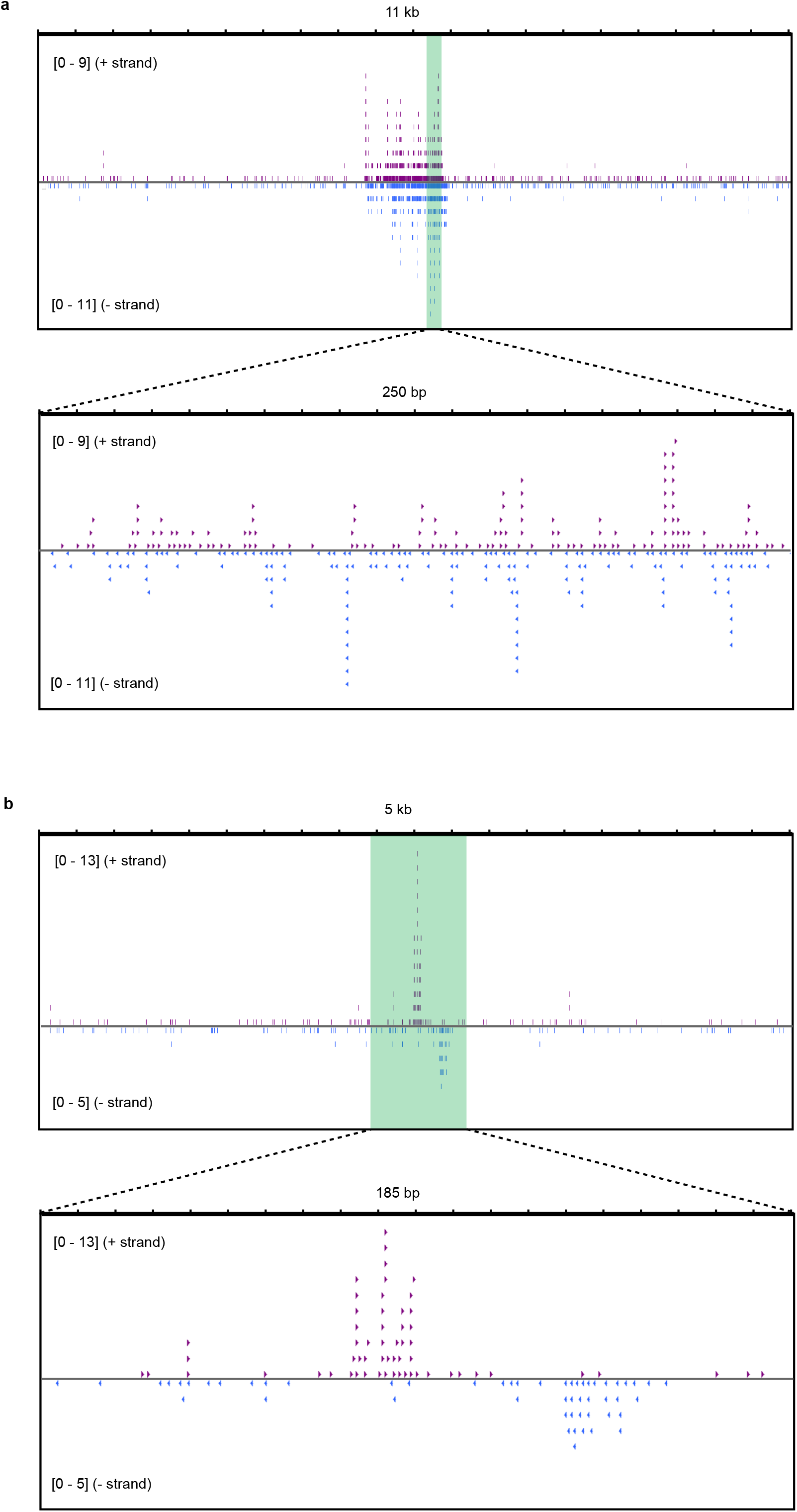
Genome browser view of two recurrent DSB positions in HEK293 cells. (**a**) 11kb view of a chr17 DSB hotspot. Purple arrows represent DSB ends labelled on the right side (+ strand) and blue arrows represent DSB ends labelled on the left side (- strand). Recurrent DSBs are evenly distributed throughout the hotspot region. (**b**) 2kb view of a chr11 DSB hotspot. Recurrent DSBs can be detected at different positions on the plus and minus strands.

**Supplementary Fig 4:**
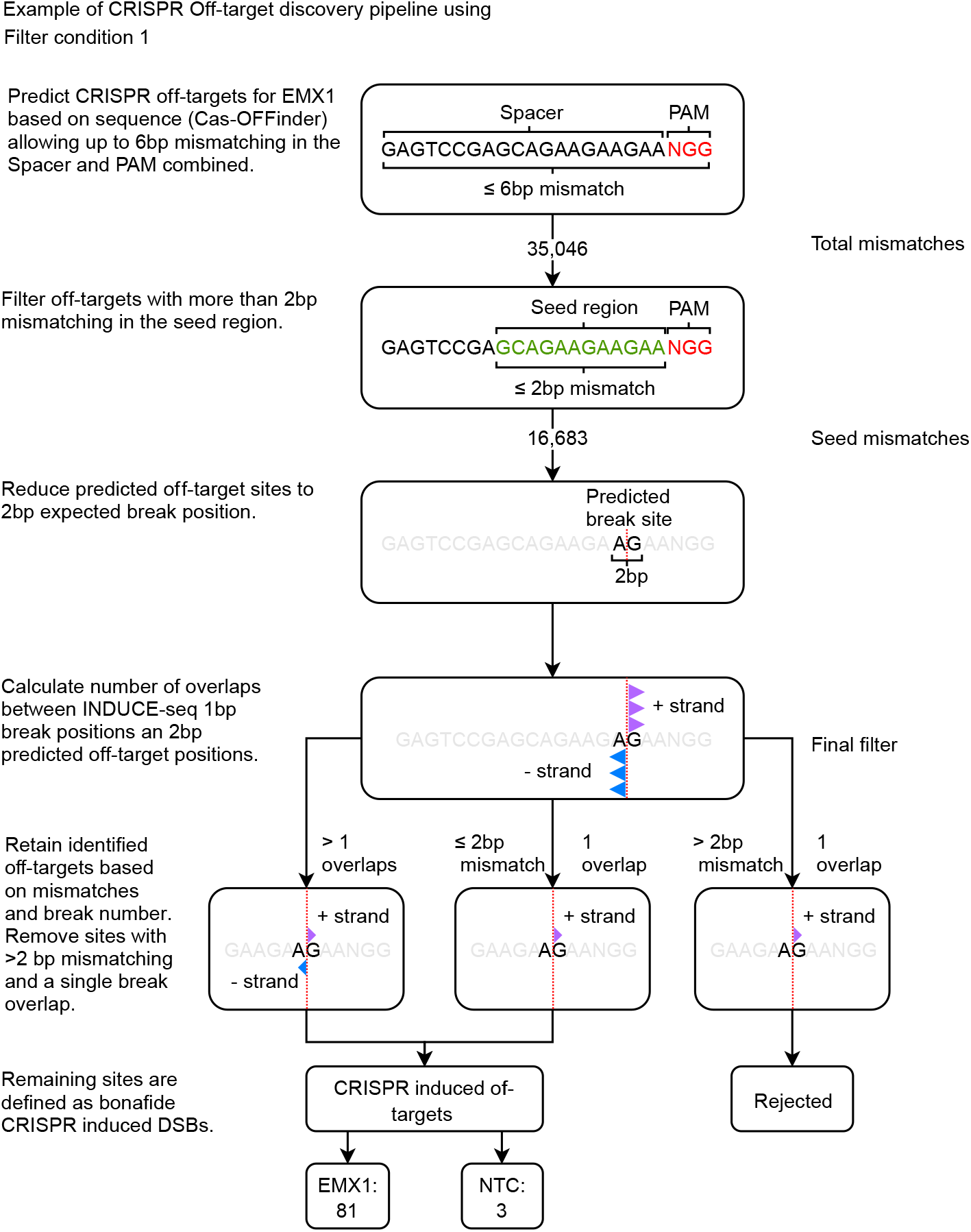
CRISPR off-target discovery pipeline. Example of the procedure for the identification of off-target CRISPR-induced DSBs. The numbers shown refer to filter condition 1 in the procedure.

**Supplementary Fig 5:**
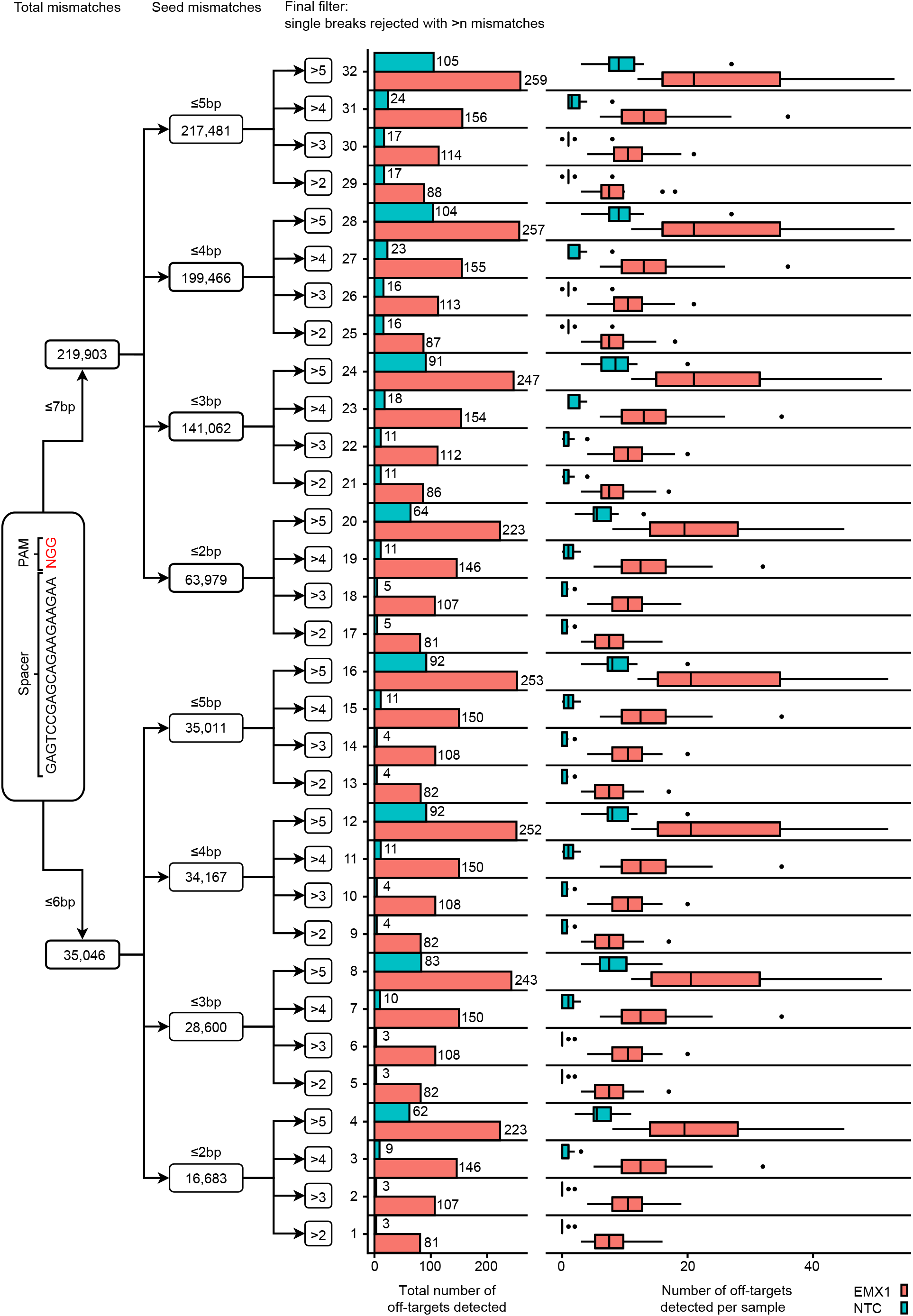
Overview of all filter conditions used for off-target discovery. (**Left side**) The three levels of filtering employed as described in the discovery pipeline. A variable number of mismatches was tested at each level of filtering, resulting in 32 combinations of filter conditions. (**Right side**) For each filter condition a set of CRISPR-induced DBSs was determined for EMX1 treated and control samples. The total number of off-target sites for all EMX1 and control samples combined is shown by the horizontal bars. The boxplots show the number off-targets detected for each individual dataset.

**Supplementary Fig 6:**
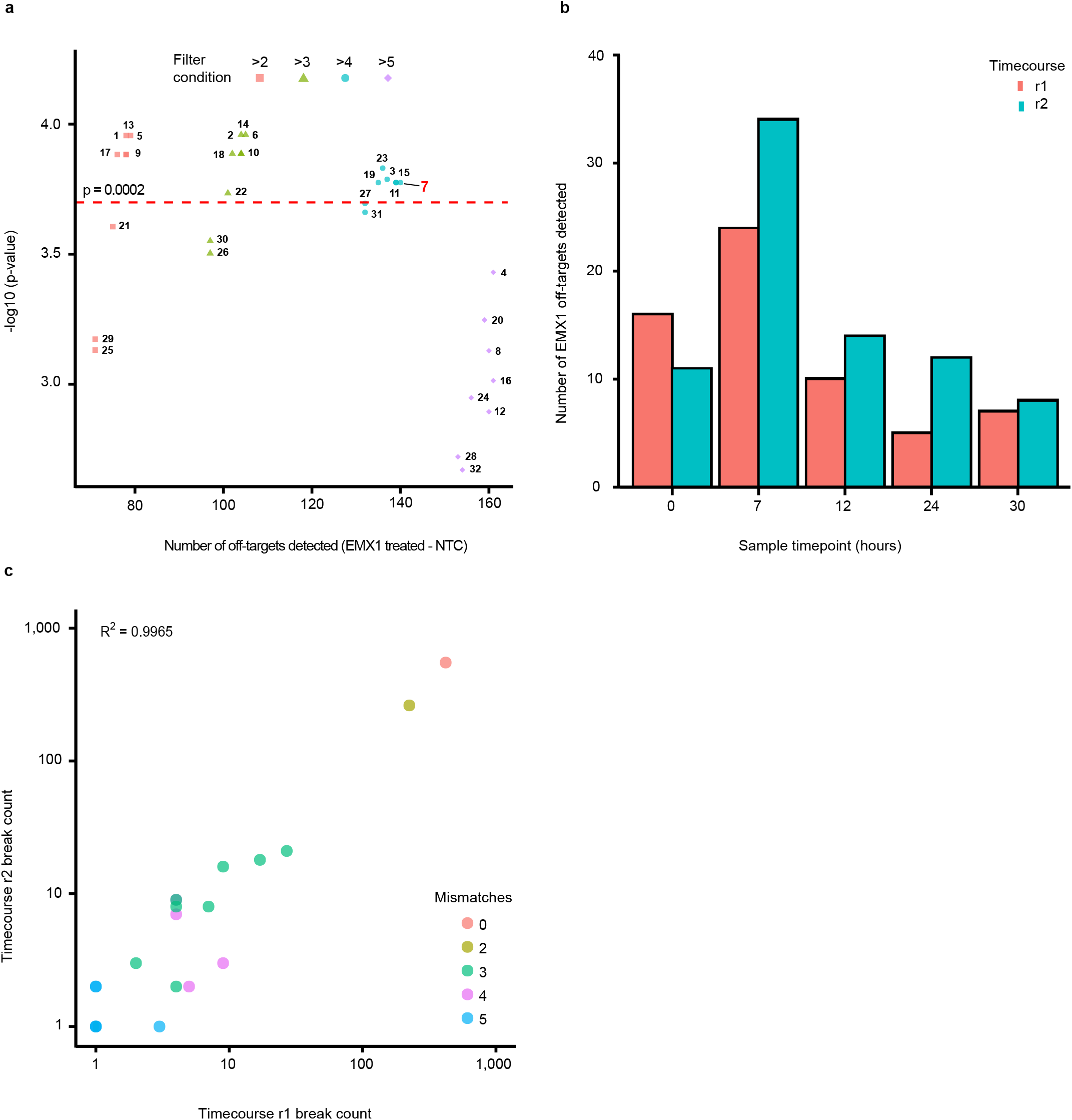
Selection of parameters for determining CRISPR off-target discovery. (**a**) Scatterplot showing the relationship between EMX1 off-target site discovery and their significance value for each filter condition (1-32). Significance level was calculated using a Wilcoxon rank sum (two-sided) test. Condition 7 (red highlight) was selected for all subsequent analysis. (**b**) Comparison between the number of EMX1 off-targets detected across r1 and r2 time course experiments. (**c**) Scatterplot showing the break number found at CRISPR off-target sites identified in both independent experiments.

**Supplementary Fig 7.**
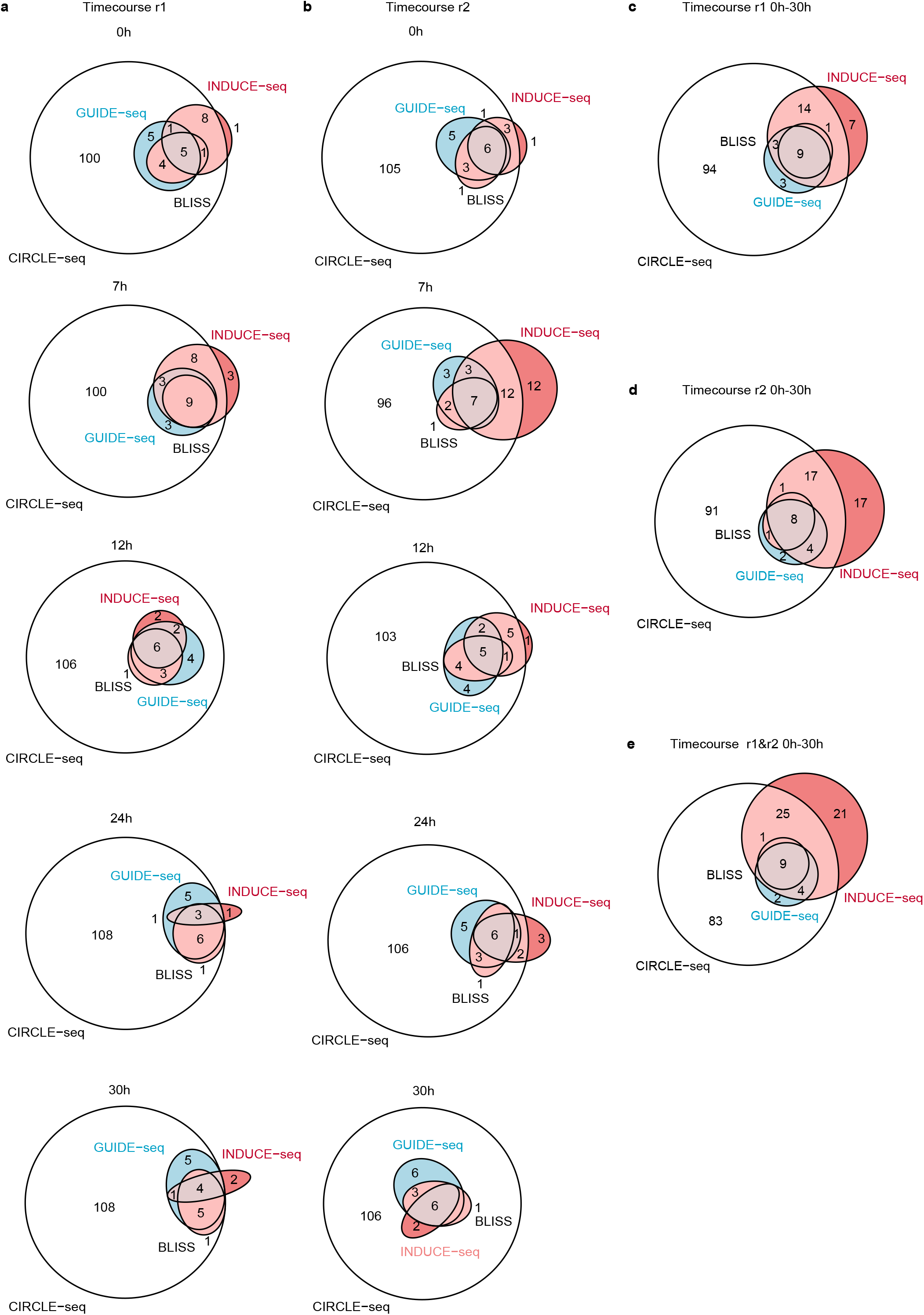
Venn diagrams showing overlaps of the off-targets identified by INDUCE-seq, CIRCLE-seq, GUIDE-seq and BLISS. (**a and b**) Overlaps calculated for samples 0h to 30h from two independent experiments r1 (**a**) and r2 (**b**). (**c and d**) The combined overlaps from all time points for set r1 (**c**) and r2 (**d**). (**e**) Overlaps calculated between methods when all INDUCE-seq samples are combined.

**Supplementary Fig 8.**
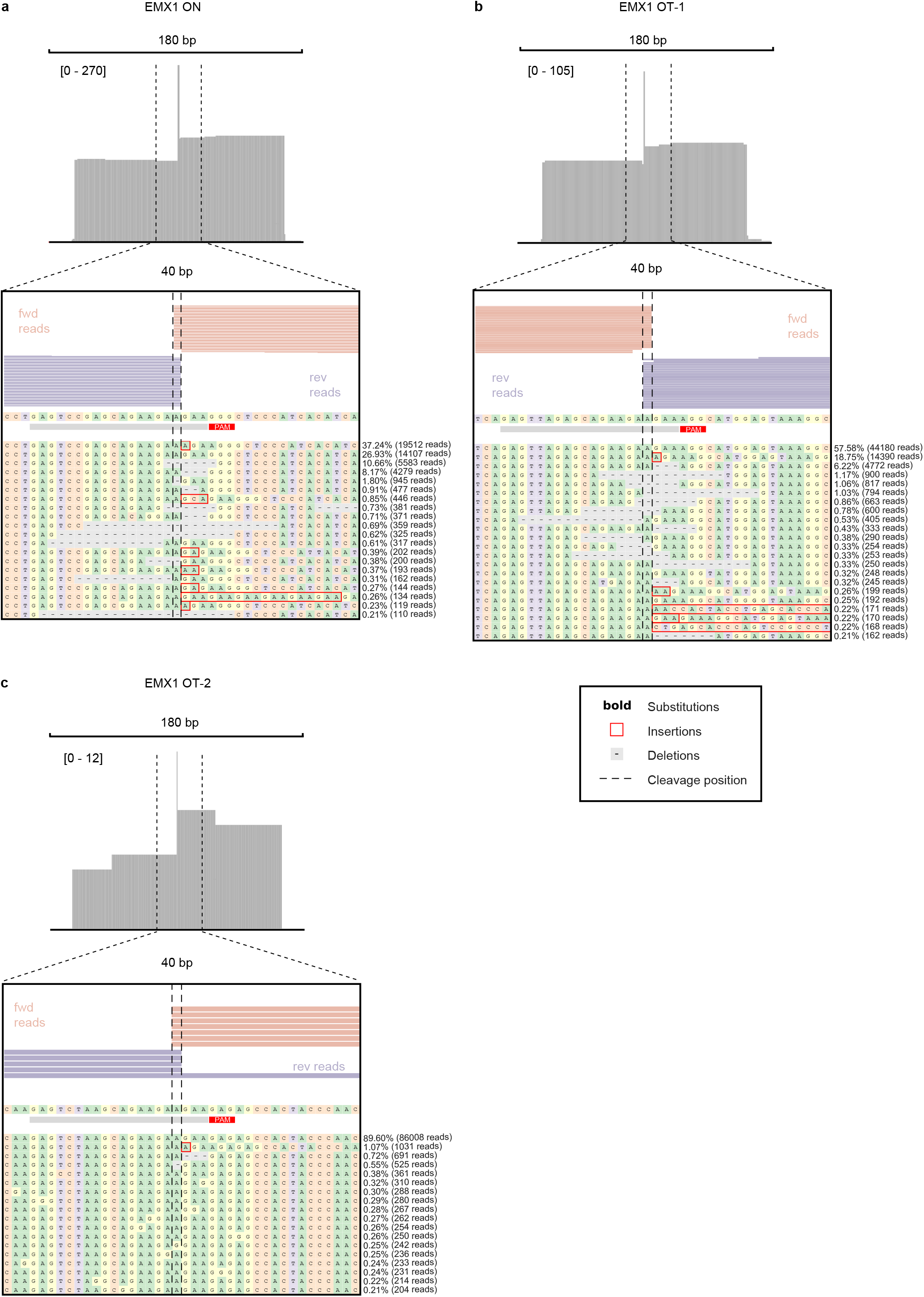
The INDUCE-seq detected DSB pattern at CRISPR induced on- and off-target sites relates to editing outcome. Coverage tracks of the EMX1 on-target (**a**) and the two top ranking off-targets, OT-1 (**b**), and OT-2 (**c**), spanning 180bp. A close-up view of the 40bp region surrounding each target site, INDUCE-seq shows a distinct 1bp overhanging cleavage pattern rather than the usual Cas9-induced blunt DSB. Corresponding indel spectra, as measured by amplicon sequencing, at each site, shows the position of the indel mutations in relation to the observed break sites.

## Online Methods

### Cell culture and treatment

HEK293, HEK293T, and U2OS DIvA cells were cultured in DMEM (Life Technologies) supplemented with 10% FBS (Life Technologies) at 37°C at 5% CO_2_. HEK293 cells were nucleofected with 224 pmol RNP per 3.5×10^5^ cells using a Lonza 4D-Nucleofector X unit with pulse code CM-130. Cells were harvested at 0, 7, 12, 24, and 30 h post nucleofection for INDUCE-seq processing. To stimulate AsiSI-dependent DSB induction, DIvA cells were treated with 300 nM 4OHT (Sigma, H7904) for 4 h.

### Cas9 protein and sgRNA

The guide RNA targeting EMX1 (GAGTCCGAGCAGAAGAAGAA) was synthesized as a full-length non-modified sgRNA oligonucleotide (Synthego). Cas9 protein was produced in-house (AstraZeneca) and contained an N-terminal 6xHN tag.

### INDUCE-seq method

Cells were seeded to 96 well plates pre-coated with Poly-D-lysine (Greiner bio-one, 655940) at a density of ~1×10^5^/well and crosslinked in 4% PFA (Pierce, 28908) for 10 min at rt. Cells were washed in 1x PBS to remove formaldehyde and stored at 4°C for up to 30 days. The INDUCE-seq method was initiated by permeabilising cells. Between incubation steps, cells were washed in 1x PBS at rt. Cells were permeabilised by incubation in Lysis buffer 1 (10 mM Tris-HCL pH 8, 10 mM NaCl, 1 mM EDTA, 0.2% Triton X-100, pH 8 at 4°C) for one hour at rt, followed by incubation in Lysis buffer 2 (10 mM Tris-HCL, 150 mM NaCl, 1 mM EDTA, 0.3% SDS, pH 8 at 25°C) for one hour at 37°C. Permeabilised cells were washed three times in 1x CutSmart^®^ Buffer (NEB, B7204S) and blunt-end repaired using NEB Quick Blunting Kit (E1201L) +100 μg/mL BSA in a final volume of 50 μL at rt for one hour. Cells were then washed three times in 1x CutSmart^®^ Buffer and A-tailed using NEBNext^®^ dA-Tailing Module (NEB, E6053L) in a final volume of 50 μL at 37°C for 30 mins. A-tailed cells were washed three times in 1x CutSmart^®^ buffer then incubated in 1x T4 DNA Ligase Buffer (NEB, B0202S) for 5 mins at rt. A-tailed ends were labelled by ligation using T4 DNA ligase (NEB, M0202M) + 0.4 μM Modified P5 adapter in a final volume of 50 μL at 16°C for 16-20 h. Following ligation, excess P5 adapter was removed by washing cells 10 times in wash buffer at rt (10 mM Tris-HCL, 2 M NaCl, 2 mM EDTA, 0.5% Triton X-100, pH 8 at 25°C), incubating for 2 mins each wash step. Cells were washed once in PBS and then once in nuclease free H_2_O (IDT, 11-05-01-04). Genomic DNA was extracted by incubating cells in DNA extraction buffer (10 mM Tris-HCL, 100 mM NaCl, 50 mM EDTA, 1.0% SDS, pH 8 at 25°C) + 1 mg/mL Proteinase K (Invitrogen, AM2584) in a final volume of 100 μL for 5 mins at rt. The cell lysates were transferred to 1.5 mL Eppendorf RNA/DNA LoBind tubes (Fisher Scientific, 13-698-792) and incubated at 65°C for 1 hour, shaking at 800rpm. DNA was purified using Genomic DNA Clean & Concentrator™-10 (Zymo Research, D4010), and eluted using 100 μL Elution Buffer. DNA yield was assessed using 1 μL sample and Qubit DNA HS Kit (Invitrogen, Q32854) before proceeding to library preparation. Genomic DNA was fragmented to 300-500bp using a Bioruptor Sonicator, and size selected using SPRI beads (GC Biotech, CNGS-0005) to remove fragments <150bp. Fragmented and size-selected DNA was end-repaired using NEBNext^®^ Ultra™ II DNA Module (NEB, E7546L). Fragmented and end-repaired DNA was added directly to the ligation reaction using NEBNext^®^ Ultra™ II Ligation Module (NEB, E7595L) according to the manufacturer’s instructions using 7.5 μM Modified half-functional P7 adapter and omitting USER enzyme addition. The ligated sequencing libraries were purified using SPRI beads. Libraries were purified twice more using SPRI beads, and size selected to remove fragments <200bp to remove residual adapter DNA. Final clean libraries were quantified by qPCR using the KAPA Library Quantification Kit for Illumina^®^ Platforms (Roche, 07960255001). Samples were pooled and concentrated to the desired volume for sequencing using a SpeedVac. Sequencing was performed on an Illumina NextSeq 500 using 1×75bp High Capacity flow cell.

### INDUCE-seq adapters

All modified INDUCE-seq adapter oligonucleotides were purchased from IDT. Single stranded oligonucleotides were annealed at a final concentration of 10 μM in Nuclease-free Duplex Buffer (IDT, 11-01-03-01) by heating to 95°C for 5 minutes and slowly cooling to 25°C using a thermocycler. INDUCE-seq P5 adapter: 5’-A*ATGATACGGCGACCACCGAGATCTACACTCTTTCCCTACACGACGCTCTTCC GATC*T-3’ and 5’-Phos-GATCGGAAGAGCGTCGTGTAGGGAAAGAGTGTAGATCT CGGTGGTCGCCGTATCATT-spacerC3-3’. INDUCE-seq half-functional P7 adapter: 5’-Phos-GATCGGAAGAGCACACGTCTGAACTCCAGTCAC-spacerC3-3’ and 5’-C*AAGCAGAAGACGGCATACGAGAT[INDEX]GTGACTGGAGTTCAGACGTGTGC TCTTCCGATC*T-3’ (*phosphorothioate bond).

### *In situ* DSB induction with HindIII

Pilot INDUCE-seq experiments were performed by inducing DSBs *in situ* in HEK293T cells using the restriction enzyme HindIII-HF^®^ (NEB, R3104S). This process was the same as described for the full INDUCE-seq method, with the addition of DSB induction prior to end blunting. Following cell permeabilization DSBs were induced using 50U HindIII-HF^®^ in 1x CutSmart^®^ Buffer in a final volume of 50 μL. Digestions were performed at 37°C for 18 hours.

### INDUCE-seq data analysis pipeline

Demultiplexed FASTQ files were obtained and passed through Trim Galore! (http://www.bioinformatics.babraham.ac.uk/projects/trim_galore/) to remove the adapter sequence at the 3’ end of reads using the default settings. Following read alignment to the human reference genome (GRCh37/hg19) using BWA-mem^24^, alignments mapped with a low alignment score (MAPQ<30) were removed using SAMtools^25^ and soft-clipped reads were filtered using a custom AWK script to ensure accurate DSB assignment. The resulting BAM files were converted into BED files using bedtools bam2bed function^26^, after which the list of read coordinates were filtered using regions of poor mappability, chromosome ends, and incomplete reference genome contigs, to remove these features from the data. DSB positions were assigned as the first 5’ nucleotide upstream of the read relative to strand orientation and were output as a ‘breakends’ BED file. Care was taken to remove optical duplicates while retaining real recurrent DSB events. By maintaining each read ID, flow cell X and Y positional information was used to filter out optical duplicates using a custom AWK script. The final output was a BED file containing a list of quantified single nucleotide break positions.

### HindIII-induced DSB analysis in HEK293T cells

The positions of HindIII target sites within hg19 were first predicted *in silico* using the tool SeqKit locate^27^, allowing a max mismatch of 2bp from the HindIII target sequence AAGCTT. The number of breaks overlapping with these predicted sites was calculated using bedtools intersect. To compare with the DSBCapture EcoRV experiment^12^, the same coverage threshold of ≥ 5 breaks per site was used to define each HindIII induced break site.

### AsiSI-induced DSB detection and analysis in DIvA cells

The positions of AsiSI target sites were calculated in the same way as for HindIII, however with no mismatches allowed and using the sequence GCGATCGC. As DIvA cells are female, sites present on the Y chromosome were removed leaving 1211 sites for chr1-X. To stringently calculate genuine AsiSI induced breaks, the 8 bp AsiSI site was reduced to 1bp genomic intervals at the predicted break positions. This reduced each 8 bp genomic interval to two 1bp intervals; at position 6 on the plus strand, and position 3 on the minus strand. Direct overlaps were then calculated between 1bp breakend positions and the predicted AsiSI break sites using bedtools intersect. Matching strand orientation was required for each overlap to be considered a genuine AsiSI-induced break site.

### CRISPR off-target analysis pipeline

Two sets of potential off-target sites for EMX1 in hg19 were first predicted using the command line version of Cas-OFFinder^28^, allowing up to 6 mismatches in the spacer and canonical PAM combined for the first set, and up to 7 mismatches for the second. Next, both sets of predicted sequences were filtered based on the mismatch number in the seed region, defined as the 12 nucleotides proximal to the PAM. Each set was filtered for up to 2, 3, 4 and 5 mismatches in the seed, generating a set of 8 files with different mismatch filtering parameters. To define CRISPR-induced DSBs, each 23 bp predicted site was first reduced to a 2bp interval flanking the expected CRISPR break position, 3bp upstream of the PAM. Overlaps were then calculated between these 2 bp expected break regions and the INDUCE-seq 1 bp breakend positions using bedtools intersect^26^, returning a set of DBSs identified at expected CRISPR break sites. Finally, DSBs overlapping with CRISPR sites were filtered based on the site mismatch number and the number of breaks detected at the site. Sites possessing mismatches >n were required to have more than 1 DSB overlap to be retained as a genuine off-target site. Each set of break overlaps was filtered using a mismatch value of >2, >3, >4 and >5, resulting in a total of 32 filter conditions and off-target datasets for each INDUCE-seq sample.

### Calculating overlaps between CRISPR off-target detection methods

EMX1 off-target sites were compared with alternative methods CIRCLE-seq, Digenome-seq, GUIDE-seq, BLISS, and HTGTS^9, 10, 18, 29, 30^. Genome interval files were generated for each respective off-target detection method. Overlaps of the EMX1 off-targets detected by each method were calculated using bedtools intersect ^26^.

### Amplicon-seq validation of mutational outcome

Amplicon sequencing DNA libraries were prepared using a custom panel of rhAmpSeq RNase-H dependent primers (IDT) that flank the INDUCE-seq identified off-targets for EMX1 (**Supplementary Data 1**). Multiplex PCR was carried out according to manufacturer’s instructions using the rhAmpSeq HotStart Master Mix 1, the custom primer mix, and 10 ng of genomic DNA. PCR products were purified using SPRI beads and Illumina sequencing P5 and P7 index sequences were incorporated through a second multiplex PCR using rhAmpSeq HotStart Master Mix 2. Resulting sequencing libraries were pooled and sequenced using an Illumina NextSeq 550 Mid Output flow cell with 2x 150bp chemistry. Editing outcomes at the on- and off-targets were determined using CRISPResso software^21^ v2.0.32 with the following parameters: CRISPRessoPooled -q30 -ignore_substitutions --max_paired_end_reads_overlap 151. Indel frequencies were compared using CRISPRessoCompare.

### Data availability

All sequencing data related to this study have been deposited in the NCBI Sequence Read Archive at PRJNA636949. All other data are available from the authors upon request.

## Acknowledgements

We thank Jessica Downs and Hugang Feng, ICR, London, UK, for their insights and for providing U2OS DIvA cells. We thank Ross Chapman, Oxford University, Oxford, UK, for his insights and reading of the manuscript. We thank Steve Jackson, Cambridge University, Cambridge, UK, for constructive discussions. We acknowledge our colleagues at the Wales Gene Park for their insight, expertise and technical support in generating the NGS data that assisted this research. Wales Gene Park is a Health and Care Research Wales funded infrastructure support group. F.M.D. is supported by a BBSRC/AstraZeneca CASE studentship (BB/P504841/1). P.V.E. is supported by the BBSRC (BB/R00756X/1).

## Author contributions

F.M.D. conceived and developed the INDUCE-seq method, conducted all experiments, performed bioinformatics analysis, interpreted results, and wrote the manuscript. P.V.E. contributed to the development of the INDUCE-seq method, contributed to the bioinformatics analysis, interpreted results, and wrote the manuscript. M.D.F. supervised the project and conceived of the study. L.L. contributed to conducting CRISPR experiments. R.N. supervised the CRISPR part of the project. S.H.R. supervised the project, conceived of the study, interpreted results, and wrote the manuscript.

## References

1. Khanna, K.K. & Jackson, S.P. DNA double-strand breaks: signaling, repair and the cancer connection. Nature Genetics 27, 247–254 (2001).

2. Vilenchik, M.M. & Knudson, A.G. Endogenous DNA double-strand breaks: Production, fidelity of repair, and induction of cancer. Proceedings of the National Academy of Sciences of the United States of America 100, 12871–12876 (2003).

3. Jackson, S.P. & Bartek, J. The DNA-damage response in human biology and disease. Nature 461, 1071–1078 (2009).

4. Chapman, J.R., Taylor, M.R.G. & Boulton, S.J. Playing the End Game: DNA Double-Strand Break Repair Pathway Choice. Molecular Cell 47, 497–510 (2012).

5. Jinek, M. et al. A Programmable Dual-RNA-Guided DNA Endonuclease in Adaptive Bacterial Immunity. Science 337, 816–821 (2012).

6. Cong, L. et al. Multiplex Genome Engineering Using CRISPR/Cas Systems. Science 339, 819–823 (2013).

7. Wienert, B. et al. Unbiased detection of CRISPR off-targets in vivo using DISCOVER-Seq. Science 364, 286–+ (2019).

8. Iacovoni, J.S. et al. High-resolution profiling of gamma H2AX around DNA double strand breaks in the mammalian genome. Embo Journal 29, 1446–1457 (2010).

9. Tsai, S.Q. et al. GUIDE-seq enables genome-wide profiling of off-target cleavage by CRISPR-Cas nucleases. Nature Biotechnology 33, 187–197 (2015).

10. Frock, R.L. et al. Genome-wide detection of DNA double-stranded breaks induced by engineered nucleases. Nature Biotechnology 33, 179–186 (2015).

11. Crosetto, N. et al. Nucleotide-resolution DNA double-strand break mapping by nextgeneration sequencing. Nature Methods 10, 361–+ (2013).

12. Lensing, S.V. et al. DSBCapture: in situ capture and sequencing of DNA breaks. Nature Methods 13, 855–+ (2016).

13. Canela, A. et al. DNA Breaks and End Resection Measured Genome-wide by End Sequencing. Molecular Cell 63, 898–911 (2016).

14. Vitelli, V. et al. Recent Advancements in DNA Damage-Transcription Crosstalk and High-Resolution Mapping of DNA Breaks. Annual Review of Genomics and Human Genetics, Vol 18 18, 87–113 (2017).

15. Aird, D. et al. Analyzing and minimizing PCR amplification bias in Illumina sequencing libraries. Genome Biology 12 (2011).

16. Jones, M.B. et al. Library preparation methodology can influence genomic and functional predictions in human microbiome research. Proceedings of the National Academy of Sciences of the United States of America 112, 14024–14029 (2015).

17. Kebschull, J.M. & Zador, A.M. Sources of PCR-induced distortions in high-throughput sequencing data sets. Nucleic Acids Research 43 (2015).

18. Yan, W.X. et al. BLISS is a versatile and quantitative method for genome-wide profiling of DNA double-strand breaks. Nature Communications 8 (2017).

19. Iannelli, F. et al. A damaged genome’s transcriptional landscape through multilayered expression profiling around in situ-mapped DNA double-strand breaks. Nature Communications 8 (2017).

20. Wang, X.L. et al. Unbiased detection of off-target cleavage by CRISPR-Cas9 and TALENs using integrase-defective lentiviral vectors. Nature Biotechnology 33, 175–178 (2015).

21. Pinello, L. et al. Analyzing CRISPR genome-editing experiments with CRISPResso. Nature Biotechnology 34, 695–697 (2016).

22. Akcakaya, P. et al. In vivo CRISPR editing with no detectable genome-wide off-target mutations. Nature 561, 416–+ (2018).

23. Vakulskas, C.A. et al. A high-fidelity Cas9 mutant delivered as a ribonucleoprotein complex enables efficient gene editing in human hematopoietic stem and progenitor cells. Nature Medicine 24, 1216–+ (2018).

24. Li, H. & Durbin, R. Fast and accurate short read alignment with Burrows-Wheeler transform. Bioinformatics 25, 1754–1760 (2009).

25. Li, H. et al. The Sequence Alignment/Map format and SAMtools. Bioinformatics 25, 2078–2079 (2009).

26. Quinlan, A.R. & Hall, I.M. BEDTools: a flexible suite of utilities for comparing genomic features. Bioinformatics 26, 841–842 (2010).

27. Shen, W., Le, S., Li, Y. & Hu, F.Q. SeqKit: A Cross-Platform and Ultrafast Toolkit for FASTA/Q File Manipulation. Plos One 11 (2016).

28. Bae, S., Park, J. & Kim, J.S. Cas-OFFinder: a fast and versatile algorithm that searches for potential off-target sites of Cas9 RNA-guided endonucleases. Bioinformatics 30, 1473–1475 (2014).

29. Tsai, S.Q. et al. CIRCLE-seq: a highly sensitive in vitro screen for genome-wide CRISPR Cas9 nuclease off-targets. Nature Methods 14, 607–+ (2017).

30. Kim, D. et al. Digenome-seq: genome-wide profiling of CRISPR-Cas9 off-target effects in human cells. Nature Methods 12, 237–+ (2015).

